# Sub-species niche specialization in the oral microbiome is associated with nasopharyngeal carcinoma risk

**DOI:** 10.1101/782417

**Authors:** Justine W. Debelius, Tingting Huang, Yonglin Cai, Alexander Ploner, Donal Barrett, Xiaoying Zhou, Xue Xiao, Yancheng Li, Jian Liao, Yuming Zheng, Guangwu Huang, Hans-Olov Adami, Yi Zeng, Zhe Zhang, Weimin Ye

## Abstract

Oral health and changes in the oral microbiome have been associated with both local and systemic cancer. Poor oral hygiene is a known risk factor for Nasopharyngeal Carcinoma (NPC), a virally-associated head and neck cancer endemic to southern China. We explored the relationship between NPC and the oral microbiome using 16s rRNA sequencing in a study of 499 NPC patients and 495 population-based age and sex frequency-matched controls from an endemic area of Southern China. We found a significant reduction in community richness in cases compared to controls. Differences in the overall microbial community structure between cases and controls could not be explained by other potential confounders; disease status explained 5 times more variation in the Unweighted UniFrac distance than the next most explanatory variable. In feature-based analyses, we identified a pair of co-excluding *Granulicatella adiacens* (*G. adicans*) amplicon sequence variants (ASVs) which were strongly associated with NPC status and differed by a single nucleotide. The *G. adicans* variant an individual carried was also associated with the overall microbial community based on beta diversity. Co-occurrence analysis suggested the two *G. adicans* ASVs sit at the center of two co-excluding clusters of closely related organisms. Our results suggest there are differences in the oral microbiome between NPC patients and healthy controls and these may be associated with both a loss of microbial diversity and niche specialization among closely related commensals.

**Importance**

The relationship between oral health and the risk of Nasopharyngeal Carcinoma has been previously established. However, the role of oral microbiome has not been evaluated in the disease in a large epidemiological study. This paper clearly establishes a difference in the oral microbiome between NPC patients and healthy controls which cannot be explained by other confounding factors. It furthermore identifies a pair of closely related co-excluding organisms associated with the disease, highlighting the importance of modern methods for single nucleotide resolution in 16s rRNA sequence characterization. To the best of our knowledge, this is one of the first examples of cancer-associated niche-specialization of the oral microbiome.

## Introduction

The microbiome, including the oral microbiome, is emerging as an important factor in cancer and carcinogenic processes. *Helicobacter pylori* is perhaps the best known example of an oncogenic bacteria; sequencing studies have implicated a new set of organisms including *Fusobacterium nucleatum* in colorectal cancer, although cancer-associated differences in the microbiome have been seen across a wide variety of tumor types (1–3). Proposed mechanisms have included genotoxicity, immune regulation, and the production of oncogenic metabolites (4). Furthermore, differences in the local microbiome have been shown in cancer-associated viral infections including chronic Hepatitis infection and Human Papilloma Virus (5–8). The microbiome is known to contribute to and modulate infection, persistence, re-activation, and transmission of oncogenic viruses, while viral infection may contribute to immune-regulation of the microbiome (9).

Nasopharyngeal carcinoma (NPC) is a virally-associated cancer endemic to areas including southern China, Southeast Asia, the Arctic, and the Middle East/North Africa. In these regions, the incidence rate of NPC is more than 20 times higher than the rest of the world (10). Infection with the Epstein-Barr virus (EBV) is the most widely accepted and extensively studied etiological factor, although its prevalence in adult population worldwide approaches 90% (11, 12). The geographic and cultural differences associated with NPC incidence suggest that both genetic susceptibility, and environmental/life-style factors such as cigarette smoking, salted fish consumption, and alcohol drinking, contribute to carcinogenesis (11, 13–15).

Several of these NPC risk factors may affect oral health and the oral microbiome (16–19). Poor oral hygiene is a risk factor for NPC, and a recent small scale study suggested differences in the oral microbiome between NPC patients and controls prior to radiation treatment (20, 21). However, it failed to fully address the way in which a NPC-related risk factor might confound this relationship; especially with regard to smoking. Therefore, we carried out a population-based case-control study in an endemic area of southern China (22). We analyzed microbial communities from 499 untreated incident NPC cases and 495 age and sex frequency-matched controls, and addressed the relationship between NPC status and the oral microbiome adjusted for potential confounders.

## Results

Study participants were recruited from the Wuzhou region in Southern China between 2010 and 2014 as part of a large population-based case-control study (22). Saliva was collected during interview. After sequencing and denoising to ASVs, there were 1066 subject’s samples which had sufficiently high-quality sequences and clinical information to be retained for analysis (Figure S1). Preliminary investigation suggested the microbiota of a small number of former smokers were highly heterogenous (n=72, 33 cases, 39 controls; Figure S2). We excluded former smokers from the final analysis, retaining 994 individuals (Table S1; Figure S1).

We aimed to address the relationship between NPC and the oral microbiome, adjusted for potential confounders. As a result, we looked for factors which might affect the oral microbiome at a community level. Our primary confounders included oral hygiene and health, tobacco use, family history of NPC, and tea consumption (14, 16, 18, 19, 23–27). We also considered a history of oropharyngeal inflammation, and the region where an individual lived and alcohol use, as covariates primarily expected to affect the microbiome, as well as salted fish consumption, which is primarily seen as a risk factor for NPC (15, 28, 29).

When comparing alpha diversity between cases and controls, we found that NPC cases showed significantly fewer overall ASVs, reduced phylogenetic diversity, and reduced Shannon diversity compared to controls (rank sum p < 0.001; Figure 1a; Table S2).; these findings did not change after adjustment for covariates which were significantly associated with alpha diversity (Figure 1b; Tables S3-S5). Hence, this suggests that patients newly diagnosed with NPC have lower overall microbial diversity than healthy controls. Our results agree with a small study of the oral microbiome in NPC patients (n=90), which also found reduced alpha diversity (21). Unlike other body sites, there is no clear relationship between salivary microbiome richness and the health of the microbial community.

**Figure 1.**
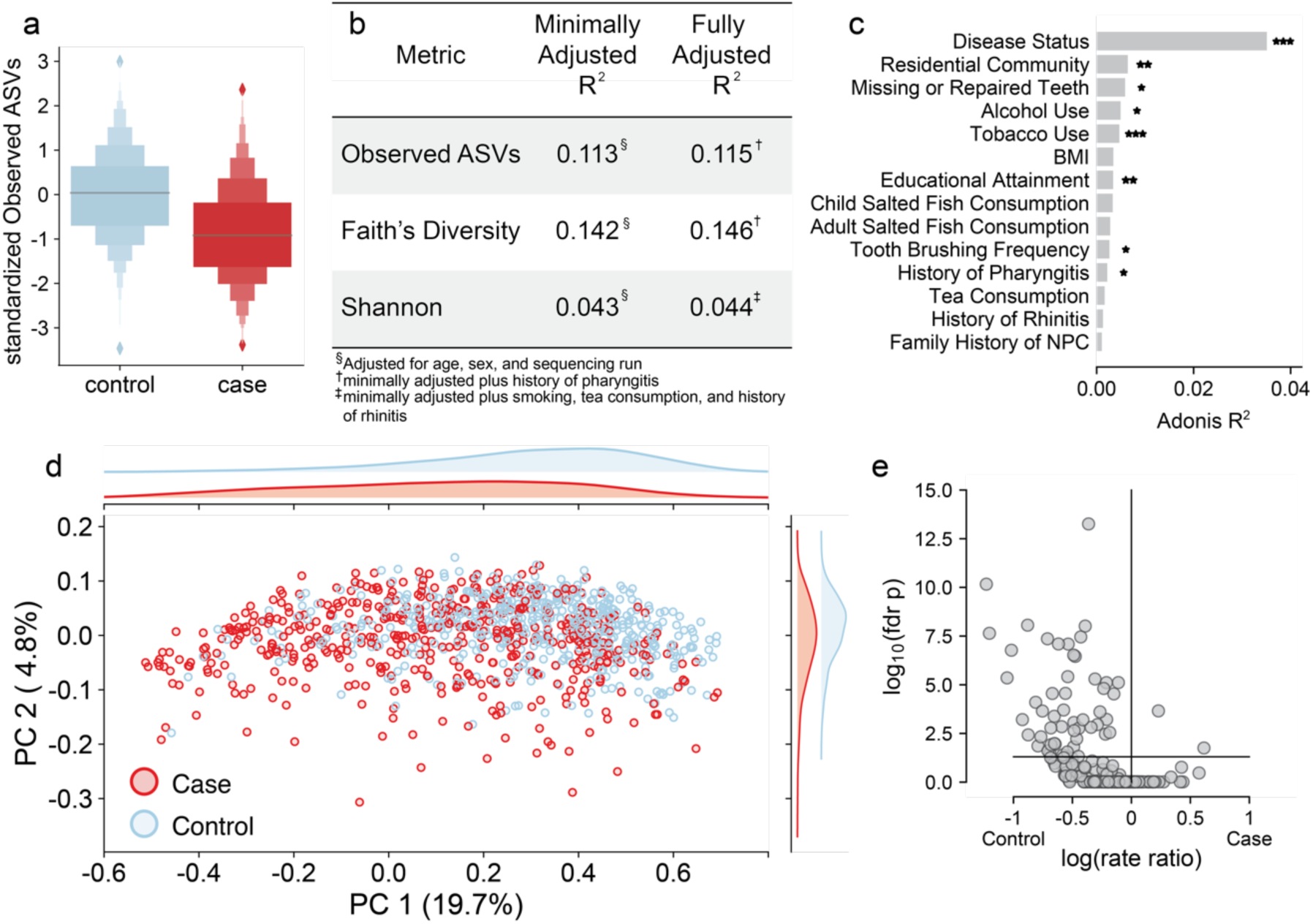
The oral microbiome differs between patients with nasopharyngeal carcinoma and healthy controls. (**a**) NPC cases (red) have significantly lower microbial richness compared to cases (blue; p < 1x10^-12^). The horizontal line in the boxlin represents the median, the large box the interquartile region, increasingly smaller boxes are the upper and lower eighths, sixteens, etc. in the data, reflecting the distribution. This difference is reflected in (**b**) the correlation coefficients from a multivariate regression model. (**c**) Adonis testing with a model adjusted for age, sex, and sequencing run shows that for unweighted UniFrac distance, NPC diagnosis has more than five times the explanatory power of the next most important variable, residential community. For 9999 permutations, FDR-adjusted p < 0.001 ***; p < 0.01 **; p < 0.05 *. (**d**) Principal coordinates analysis (PCoA) of unweighted UniFrac shows separation between cases (red) and controls (blue) along PC1 and PC2. Upper and right panels reflect the density distribution along each axis. The axes are labeled with the variation they explain. In unweighted UniFrac, PC1 explains 19.7% and PC2 explains 4.8% of the variation. A volcano plot of (**e**) the Poisson regression coefficient for disease status vs the log p-value reflects reduced diversity. The horizontal line indicates significant at a Benjamini-Hochberg corrected p-value of less than 0.05.

Similarly, when comparing global community patterns (beta-diversity) via Adonis models minimally adjusted for sex, age and sequencing run, we found significant differences between NPC cases and controls, both based on unweighted UniFrac distance as well as for weighted UniFrac and Bray-Curtis distances (FDR p< 0.001, 9999 permutations; Figures 1c,d and S3a,b) (30–32). Compared to the potential confounders in the same setting, NPC status was the strongest explanatory factor for unweighted UniFrac distance, more than five times the effect size of the next strongest variable, as well as the second-strongest factor for weighted UniFrac- and Bray-Curtis distances, just after tobacco use. There was no statistically significant difference in dispersion between cases and controls in any metric, supporting the idea that the differences are due to consistent differences between cases and controls (p > 0.55, 999 permutations; Figure 1d). Significance persisted in more fully adjusted Adonis models including potential confounders with robust differences in community patterns.

These findings establish that NPC status and smoking are strongly associated with differences in the oral microbiome in our population; the association with NPC is especially strong with regard to presence and absence of organisms (as emphasized by unweighted UniFrac), but second only to smoking with regard to abundances (as captured by weighted UniFrac and Bray-Curtis). We found no evidence that these associations are driven by community heterogeneity; they are, however, robust under adjustment for observed confounders, and in the case of the unweighted UniFrac distances, unlikely to be the result of confounding by unobserved factors due to the crushing dominance of the signal for NPC status. Since we recruited incident, treatment-naive patients, it is also implausible that the observed differences in microbiome composition are treatment-related (21, 22). Taken together, our findings provide strong evidence for a clear difference in the oral microbiome between patients with NPC and healthy controls.

Since the relationship between the microbiome and NPC status was strongest in unweighted UniFrac distance, which focuses on presence and absence, we evaluated the relationship between ASV prevalence and disease in a fully adjusted log binomial model. To limit spurious correlations, we defined presence as a relative abundance greater than 0.02% and focused on ASVs present in at least 10% of samples (n=245, Figure S4). We identified 53 ASVs which were significantly different between cases and controls (FDR p < 0.05; Figure 1e; Table S6). The large majority of these ASVs were more prevalent in controls and came from a wide variety of taxonomic clades, which may suggest a somewhat stochastic loss of ASVs in NPC patients, rather than a systematic loss of specific organisms (Table S6). This finding is in line with our alpha diversity findings, and may indicate overall community instability. In contrast, two ASVs were more prevalent in cases: a member of genus *Lactobacillus* (Lact-eca9) and a *Granulicatella* ASV (Gran-7770).

To evaluate whether NPC status affected abundance-based partitioning of the microbial community, we applied Phylofactor (33). Our model looked for phylogenetic clades which differentiated NPC cases and controls, adjusting for potential confounders (Figure 2, Table S7). Of the twelve factors examined, nine were associated with disease status. The primary partition in the data suggested a *Granulicatella* ASV (Gran-7770) was 3.4 (95% CI 2.4, 4.9) fold more abundant in NPC cases compared to controls. The third factor identified was second *Granulicatella* ASV (Gran-5a37) as less abundant in cases. Both ASVs were also associated with smoking status. We identified three large-scale shifts in microbial abundance associated with NPC status. The remaining factors associated with NPC status were all single ASVs which differentiated cases and controls, none of which differed in prevalence (Table S6, S7).

**Figure 2.**
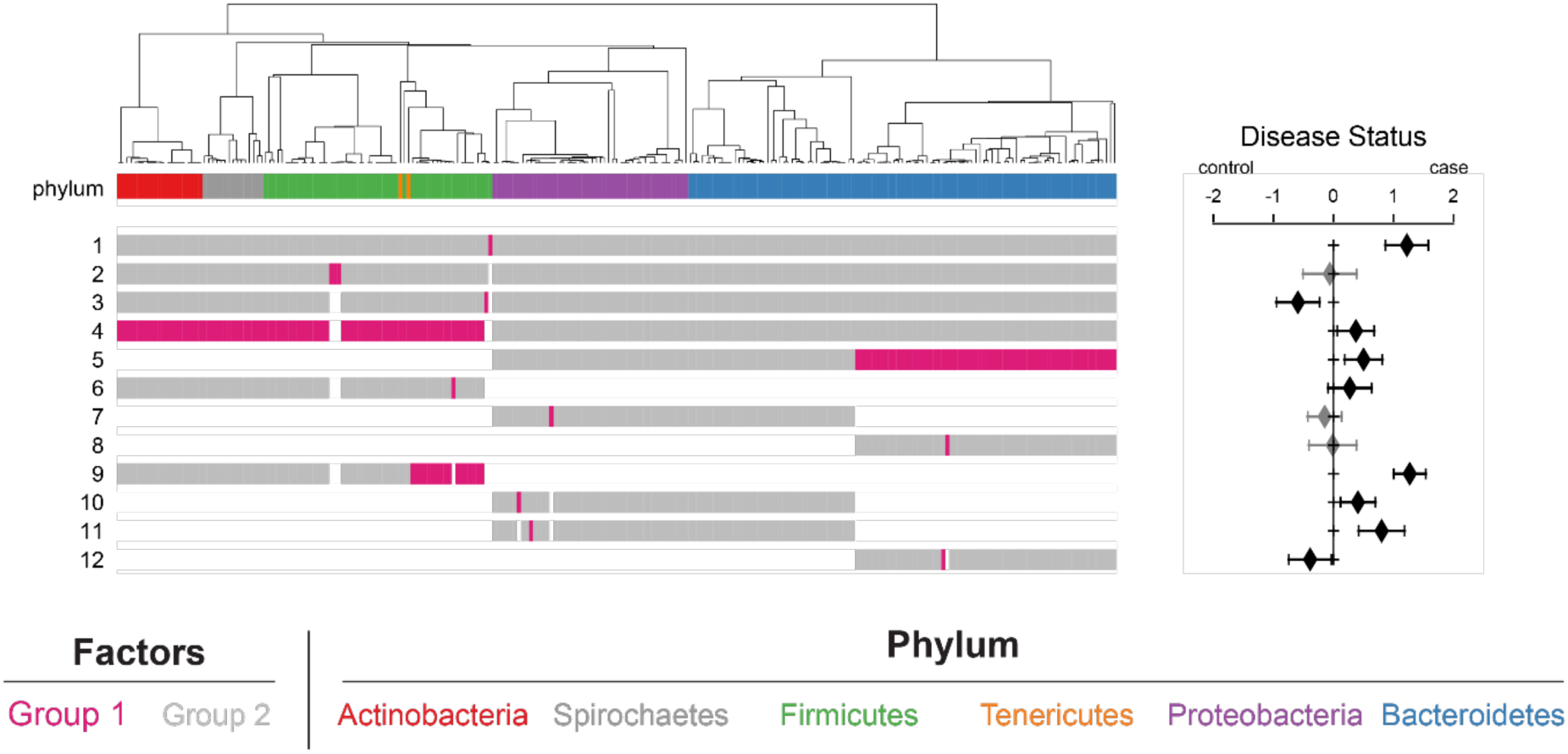
There are significant associations between phylogenetic partitioning of the taxa and NPC status. The phylogenetic tree with the first 12 phylofactor-based clade partitions is shown on the left. The top row is colored by phylum, the associated color is shown below. The isometric log transformation is taken as the ratio of the tips highlighted in pink over those highlighted in gray and passed into the regression model to predict the coefficient shown in the forest plot. Clades which are excluded from that factor appear white in the row. The forest plot to the right shows the estimated increase in the factor associated with case control status based on fitting the ratio in a linear regression adjusted for age, sex, sequencing run, number of missing or repaired teeth, tobacco use, and residential community. Error bars are 95% confidence interval for the regression coefficient. Black bars indicate significance at *a* < 0.05, gray indicates a non-significant association.

Based on the significant difference in abundance and prevalence of ASVs from genus *Granulicatella* between cases and controls, we further explored this genus. We identified a total of 14 ASVs in the dataset; three were prevalent enough to be included in our feature-based analyses (Gran-5a37, Gran-7770, and Gran-6959). In 972 (97.8%) individuals, the abundant ASVs were the only *Granulicatella* present. When blasted against the Human Oral Microbiome Database (HOMD), the ASV sequences mapped to two cultured species with more than 99.5% accuracy: *Granulicatella elegans* (*G. elegans*) which included Gran-6959 and *Granulicatella adiacens* (*G. adiacens*; Gran-7770 and Gran-5a37) (34). Strikingly, we found our two abundant *G. adiacens* ASVs differ by a single nucleotide: Gran-7770 carries a G at nucleotide 119 of our sequence (corresponding approximately to 458 in the full length 16s rRNA sequence) while Gran-5a37 carries an A.

Gran-7770 was found to be 26% more prevalent among cases, while Gran-5a37 was among the 51 ASVs less prevalent in cases (PR 0.81 [95% CI 0.74, 0.88]; Table S6]). Both ASVs were also significantly associated with smoking status: Gran-7770 was more prevalent in smokers (PR 1.48, [95% CI 1.29, 1.70]) and Gran-5a37 less prevalent (PR 0.74, [95% CI 0.67, 0.81]). There was not a significant relationship between Gran-6959 (*G. elegans*) and either disease status (PR 0.94 [95% CI 0.88, 1.00]) or tobacco use (PR 0.97 [95% CI 0.90, 1.06]).

We found that 993 out of 994 individuals carried at least one *G. adiacens* with a relative abundance of at least 0.02%: 330 (33.2%) carried only Gran-5a37, 316 (31.8%) carried Gran-7770 alone, and 347 (34.9%) carried both. Among individuals who were classified as carrying only one ASV (Gran-7770 alone or Gran 5a37 alone), the “present” ASV was at least 50-fold more abundant than the other variant. We used a multinomial logistic regression to confirm that disease status was significantly associated with variants an individual carried: compared to the odds of carrying Gran-5a37 alone, cases had significantly higher odds of carrying both ASVs and, again, significantly higher odds of carrying Gran-7770 alone (Figure 3a). Although smokers were more likely to have both ASVs or Gran-7770 alone, there was no significant interaction between smoking and disease status.

We also investigated how the presence of a *G. adiacens* variant structured the overall microbial community. We filtered the full ASV table to remove any *Granulicatella* ASVs and used the reduced table to re-calculate beta diversity metrics. The *Granulicatella*-free community recapitulated the patterns seen in the full community well (Mantel R^2^> 0.91; p=0.001, 999 permutations). We found significant differences between individuals who carried Gran-7770, both, or Gran-5a37 in weighted and unweighted UniFrac distances and Bray Curtis; all three metrics show clear separation in PCoA space (p=0.001, 999 permutations; Figure 3b; Figure S5). In unweighted UniFrac space (Figure 3b), the separation was primarily along PC2, likely corresponding to the separation along PC2 seen between cases and controls (Figure 1d). Furthermore, we found that the *G. adiacens* variant explained 16% of the variation attributed to case-control status in unweighted UniFrac distance and 15% of the variation in weighted UniFrac distance. Our results suggest that the *G. adiacens* variant carried by an individual is significantly associated with community structure, and may be a route by which NPC status shapes the oral microbiome.

**Figure 3.**
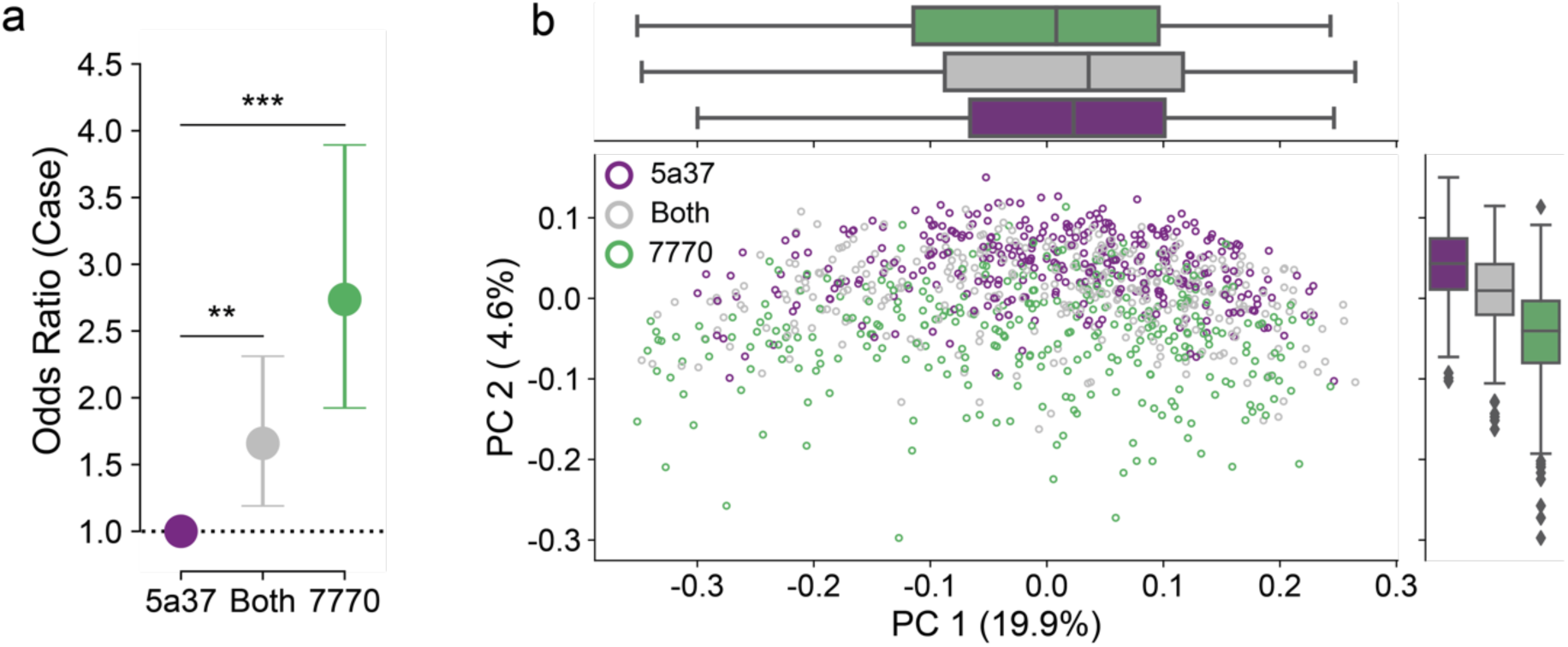
The *Granulicatella adiacens* variant predicts community structure. (**a**) NPC cases have significantly higher odds of carrying both Gran-5a37 and Gran-7770 than Gran-5a37 alone, and again, significantly higher odds than carrying either Gran-5a37 and Gran-7770 or Gran-7770. (**b**) In unweighted UniFrac space, we see separation based on the G. adiacens variant along PC2.

We used a SparCC-based network analysis to identify other community members *Granulicatella* might interact with to exert an effect on the microbiome (35). We were able to identify five networks: one pair of co-occurring ASVs, two pairs of co-excluding ASVs, one three-member network of co-occurring ASVs and a large 29-member network of co-occurring and co-excluding ASVs (Figures 4a). This main network consisted of two clusters of a total of 20 organisms which were positively correlated with a *Granulicatella* variant; the main members of the networks belonged to *Veillonella*, *Streptococcus*, and *Prevotella*. Blasting against HOMD, we identified two additional pairs of ASVs that co-excluded between the two nodes but mapped to the same clones: Stre-900d and Stre-0531 (*Streptococcus parasanguinis* clade 411) and *Prevotella melaninogenica* (Prev-b7f2 and Prev-71e7; Figure 4b; Table S8) (34).

**Figure 4.**
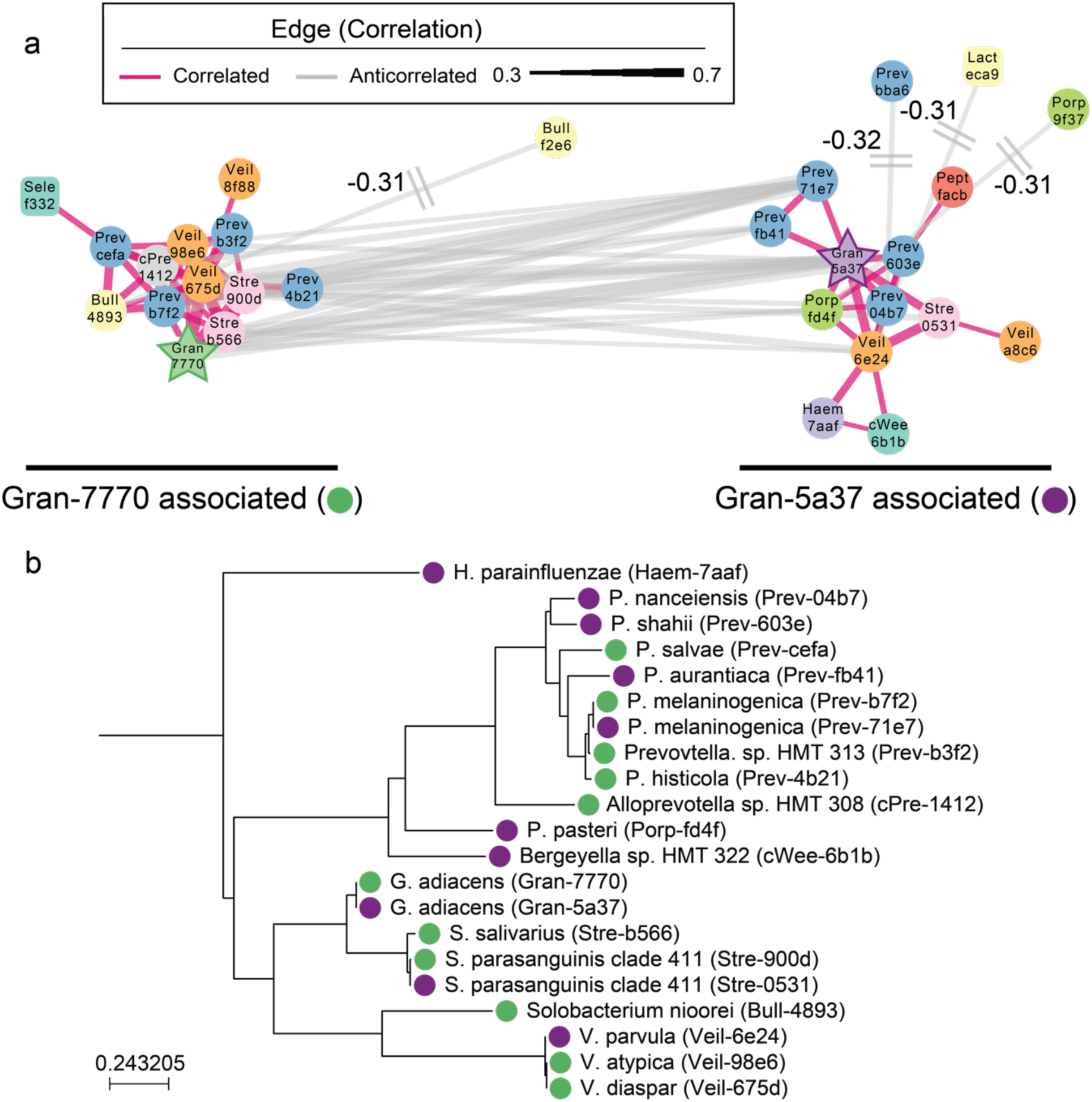
*Granulicatella adiacens* variants set at the center of a network of closely related co-occurring organisms. **(a)** SparCC-based network analysis for co-occurring and co-excluding ASVs for all subjects showed a large network with two clusters with common core structures. The color and shape of the nodes are genus-specific. The two *G. adiacens* variants are highlighted as stars: Gran-5a37 in purple and Gran-7770 in green. Correlated edges are shown in pink, anti-correlated edges are grey. The sides of each network are labeled with their associated *G. adiacens* variant. **(b)** Phylogenetic tree of the core ASVs from the network (positively correlated with either Gran-7770 or Gran-5a37). Tips are labeled by their association with Gran-7770 (Green) or Gran-5a37 (Purple).

We hypothesize the co-excluding networks of ASVs, centered around *Granulicatella*, may reflect partial niche specialization. Previous work suggests quorum sensing networks can form between the core species, and that metabolic changes occur in these networks (36, 37). These closely related organisms occupy the same niches within these metabolic networks, however, strain-specific variation may either respond to or promote disease-associated transformation. Without culture-based experimentation, it is difficult to determine how these organisms may function in concert. One major challenge for in-silico validation is the limited resolution of existing databases; our results exceed the OTU-based resolution used in construction and span a less frequently characterized hypervariable region.

Within the context of NPC in an endemic region, we hypothesize the oral microbiome may act through several potential mechanisms. The oral microbiome has been suggested to contribute to local tumorigenesis through immune regulation or oncogenic metabolites such as acetaldehyde or nitrosamines (38). An *in silico* study suggested that commercially available strains of *G. adiacens* and co-abundant organisms encode genes involved in nitrate and nitrite reduction (39). In summary, we have demonstrated a difference in the oral microbial community between NPC patients and healthy controls in an endemic area of southern China, which cannot be explained by other measured factors. The difference is associated with both a loss of community richness and differences among specific organisms, including closely related ASVs from genus *Granulicatella*. In addition, we identified a network of co-occurring and co-excluding ASVs which included these *Granulicatella* variants. These results strongly suggest a relationship between the oral microbiome and nasopharyngeal carcinoma status in untreated patients.

## Acknowledgements

The authors which to thank the study participants, the field work team for the NPCGEE project, and the Wuzhou Health System Key Laboratory for Nasopharyngeal Carcinoma Etiology and Molecular Mechanism and the Key Laboratory of High-Incidence-Tumor Prevention & Treatment (Guangxi Medical University), especially Suhua Zhong, Xiling Xiao, for the processing of salivary samples. The data was stored in the Department of Medical Epidemiology and Biostatistics; the authors wish to thank the IT group for their assistance.

We acknowledge funding by the National Cancer Institute of the NIH grant under Award Number R01CA115873 [Principal Investigator (PI): H.-O. Adami; co-PIs: Y. Zeng, Y.X. Zeng]; from the Swedish Research Council (2015-02625, 2015-06268, 2017-05814 to Ye); the National Natural Science Foundation of China (81272983 to Z. Zhang); and the Guangxi Natural Science Foundation (2013GXNSFGA019002 to Z. Zhang). T. Huang is partly supported by a grant from China Scholarship Council (201408450018).

## Data Availability

Raw sequencing data, feature table, and metadata are available from the corresponding author upon request.

## Author contributions

The study approach was conceived by GH, HA, WY, YZ, and ZZ. AP, DB, JWD, TH, WY, and YC, refined the study design for this project. JL, YC, YL, and YZ were responsible for sample collection and management. DB, SZ, and XX performed the lab work, supervised by TH, XX, XZ and ZZ. AP, JWD, and TH and performed statistical modeling and refinement; bioinformatic and biostatistical analyses were performed by JWD. WY contributed to the supervision and coordination of the project. JWD and TH wrote the manuscript; AP provided critical edits. All authors reviewed and approved the final submission.

## Methods

### Survey metadata and sample collection

Participant recruitment has been previously described (22). Briefly, incident cases of NPC in the Guangxi Autonomous Region between 2010 and 2013 were invited to participate in the full study. Age and sex matched controls were selected from the total population. The study was approved by the Institutional Review Board or Ethical Review Board at all participating centers. All study participants provided written or oral informed consent.

A questionnaire covering demographics, diet, residential, occupational, medical and family history was administered in a structured interview. Sample collection occurred at the interview. Participants were asked not to eat nor chew gum for 30 minutes prior to sample collection. Saliva samples (2ml-4ml) were collected into 50ml falcon tubes with a Tris-EDTA buffer. Demographic characteristics of the study population were compared using a two-sided t-test for continuous covariates (age) and a chi-squared test for categorical covariates. Tests were conducted using scipy 0.19.1 in python 3.5.5 (40).

### DNA extraction, PCR, and sequencing

Saliva DNA was extracted using a two-step protocol including the sample pre-processing with lysozyme lysis and bead beating, and the TIANamp blood DNA kit (Beijing, China). The 16s rRNA amplicon library was amplified with 341F/805R primers (CCTACGGGNGGCWGCAG, GACTACHVGGGTATCTAATCC) (41, 42). Samples were amplified with 20 cycles of a program with 30 seconds at 98°C for melting, 30 second at 60°C, and 30 seconds at 72°C. Samples were barcoded in a second PCR step (41). DNA clean-up was performed using Agentcourt AMPure XP purification kit. DNA volume and purity were measured on an Agilent 2100 Bioanalyzer system and Real-time polymerase chain reaction. Sequencing was performed at Beijing Genome Institute (BGI) on an Illumina MiSeq using a 2x300bp paired end strategy.

### Denoising, Annotation and Filtering

Samples were demultiplexed using an in-house script. Adaptors were trimmed and paired end sequences were joined using VSEARCH (v. 2.7) (43). Paired sequences were loaded into the November 2018 release of QIIME 2 (44). Sequences were quality filtered (q2-quality-filter) and denoised using deblur (v. 1.0.4; q2-deblur) with the default parameters on 420 bp amplicons to generate amplicon sequence variants (ASVs) (45, 46). A phylogenetic tree was built using fragment insertion into the August 2013 Greengenes 99% identity tree backbone with q2-fragment-insertion; taxonomic assignments were made with a naïve Bayesian classifier trained against the same reference (q2-feature-classifier) (47–49). In cases where the classifier or reference database was unable to describe a taxonomic level (for instance, a missing genus), the taxonomy was described by inheriting the lowest defined level using a custom python script. Following sequencing and denoising, 24,763,933 high quality reads were retained.

Any sample with fewer than 1000 reads after denoising was excluded, leaving 1074 saliva samples and 9 negative or single organism controls. Additionally, samples missing information on tobacco use, defined information about tooth brushing frequency, or an undefined residential region (n=8) were excluded (Figure S1).

Preliminary investigation suggested that the microbial communities for former smokers (*n*=72) were highly heterogenous (Figure S2). Sensitivity analyses suggest their exclusion does not alter the major community-level differences. Therefore, they were excluded, leaving a total of 994 individuals in the analysis.

ASV-based analyses were performed on a representative subset: those with at least 0.02% relative abundance in at least 10% of samples (n=245). A Mantel test was applied to Bray Curtis distance showed a correlation of 0.96 between the filtered matrix rarefied to 5000 sequences/sample and the full table distance matrix (p=0.001, 999 permutations); the mantel corresponding correlation for UniFrac distance was 0.76 (p=0.001, 999 permutations; Figure S3) (30, 32, 50).

The sequences and identifiers for the abundant ASVs are listed in supplemental file 2. ASVs are identified by the first 4 letters of their lowest taxonomic assignment and the first 4 characters of a MD5 hash of the sequence. The full taxonomic assignment and MD5 hashes can be found in Table S6.

### Diversity Analyses

Diversity analyses were performed using samples rarefied to 6,500 sequences.

Alpha diversity was calculated as observed ASVs, Shannon diversity, and Faith’s phylogenetic diversity using q2-diversity in QIIME 2 (51, 52). Potentially significant alpha diversity categories predictors were identified using a rank-sum test in scipy 0.19.1 (40). A p-value of 0.05 was considered the threshold for borderline significance for inclusion in a subsequent regression model. Alpha diversity was then evaluated in a multivariate ordinary least squares (OLS) regression model adjusted for age, sex and sequencing run number. A final model for each metric was selected by forward selection using models which resulted in decreasing AIC. We checked for the normality of residuals by plotting. The relative contribution of each covariate to that metric was estimated by a “leave one out” approach. Regressions were performed in Statsmodels (v. 0.9.0) (53). For visualization, we calculated z-normalized alpha diversity using the mean and standard deviation in diversity for the controls. Alpha diversity was plotted using boxenplots in Seaborn 0.9.0 (54, 55).

Beta diversity was measured using the unweighted UniFrac, weighted UniFrac, and Bray-Curtis metrics on rarefied data (q2-diversity) (30–32). Beta diversity was compared using Adonis in the R *vegan* library (v 2.5-2) adjusted for host age, sex, and sequencing run, with 9999 permutations (56–58). We used a permdisp test with 999 permutations and the centroid estimate to test for the presence of differences in within-group variation implemented is scikit-bio 0.5.4 (www.scikit-bio.org).(59) Uncorrected p-values of less than 0.05 were considered to have significant dispersion, since we were more concerned about false positives than false negatives. Principal coordinate analyses (PCoA)s were visualized using Emperor (v. 1.0.0b18) and seaborn v. 0.9.0 in matplotlib v. 2.2.3 (54, 60).

### ASV regression model

To look at the relationship between ASV prevalence and disease and smoking status, we used a log binomial regression which was approximated via a Poisson regression with robust standard errors, implemented via base function *glm* in R and the robust error mechanism implemented via packages *lmtest* (v 0.9) and *sandwich* (v. 2.5) in R 3.5 (58, 61–63). The model was adjusted for age, sex, sequencing run, residential community, and the number of missing or repaired teeth. “Presence” was defined as a relative abundance of 1 / 5000, which corresponded to the shallowest sequencing depth for the abundant counts. ASVs which were present in more than 1000 samples were excluded from prevalence analysis. A Benjamini-Hochberg FDR corrected p-value of 0.05 was considered significant.

### Phylofactor

Phylofactor (v. 0.01) was used to look at the relationship between disease status and phylogenetic partitioning between clades (33). Phylofactor is a compositionally aware technique which uses isometric log transforms over an unrooted phylogenetic tree to model differences in the data. This allows the partitioning of data into polyphyletic clades. The Phylofator multivariate model for each partition was modeled with an OLS regression considering diagnosis, adjusted for residential community, age, sex, number of missing or repaired teeth, tobacco use, and sequencing run. We looked at the first 12 factors using the default parameters, which optimized for explaining maximal variance. The cladogram, and regression coefficient plots were generated in seaborn (54).

### Granulicatella

Total *Granulicatella* was identified by filtering the full ASV table for any ASV assigned to the genus. Species-level assignments were made by blasting each ASV against the Human Oral Microbiome Database using the online tool; species-level assignments were taken for the cultured species with the best match (34). We treated the abundance of Gran-6959 as the *G. elegens* abundance and the combined abundance of Gran-5a37 and Gran-7770 as the *G. adiacens* abundance throughout. One sample which did not contain Gran-7770 or Gran-5a37 was excluded from *G. adicans* associated analyses.

We used a multinomial logistic regression model, implemented in the *nnet* library (v. 0.8) in R to look at whether the carriage of Gran-5a377 alone, Gran-7770 alone, or both ASVs was associated with smoking and disease status (64). The regression was adjusted for age, sex, sequencing run, number of missing or repaired teeth, residential community, the relative abundance of *G. adiacens*, and the relative abundance of *G. elegens*. Having Gran-5a37 was considered the reference group for the multinomial regression.

The effect of *Granulicatella* on alpha and beta diversity was calculated by first filtering out all Granulicatella ASVs from the table, and then rarifying to 6250 sequences/sample before diversity calculations. Adonis coefficients were calculated in a model adjusted for *G. adiacens* abundance, sequencing run, age, sex, residential community, number of missing or repaired teeth, tobacco use, and disease status. The proportion of disease status explained by comparing a model excluding the Granulicatella variant minus the model including the variant over the model excluding the variant.

### Network Analysis

We used the Sparse Cooccurrence Network Investigation for Compositional data (SCNIC; https://github.com/shafferm/SCNIC) in QIIME 2 (q2-SCNIC) to perform network analysis on the abundant ASVs. The correlation network was built using SparCC, and the network was built using edges with a correlation co-efficient of at least 0.3, allowing both co-occurrence and co-exclusion (35). Network clusters were identified by finding the most connected node and following all positively correlated nodes in the trimmed SparCC network. Networks were visualized in Cytoscape (v. 3.7.1) using a perfuse-weighted network layout (65). Nodes which were anti-correlated with a single node in the main cluster were trimmed for the sake of visualization; these are labeled with the correlation coefficient.

The phylogenetic tree of core network members was visualized using ete3 (v. 3.1.1) in python 3.6 (66).

## Notes

#### Summary of Updates

Updated text and title to focus more broadly on NPC as a viral cancer, rather than simply NPC as a rare cancer

