## Supplemental File 1 for "Sub-species niche specialization in the oral microbiome is associated with nasopharyngeal carcinoma risk"

*Supplemental Tables*

Table S1. Demographic characteristics of the study population

Table S2. Univariate association between patient characteristics and alpha diversity

Table S3. Model selection for Regression against Faith's Phylogenetic Diversity

Table S4. Model selection for Observed ASVs

Table S5. Model selection for regression against Shannon diversity

Table S6. Poisson regression coefficients

Table S7. Phylofactor Partitions

Table S8. Species-level identification of core network-related organisms

*Supplemental Figures*

Figure S1. Flowchart for sample selection

Figure S2. Former smokers have highly heterogenous microbiomes.

Figure S3. A community of abundant ASVs retains the difference in the microbiome between NPC cases and controls.

Figure S4. The abundant ASVs recapitulate community structure well.

Figure S5. *Granulicatella adiacens* variant predicts community structure in weighted metrics

**Supplemental File 2. Representative sequences for abundant ASVs**

**Supplemental File 3. Table S6**

**Supplemental File 4. Table S7**

**Table S1. Demographic characteristics of study population**

| | Cases (n=499) | | Controls (n=495) | | p-value§ |
|---|---|---|---|---|---|
| Mean age in years | 48.4 | (10.5) | 49.5 | (10.4) | 0.09† |
| Number male (%) | 356 | (71.3%) | 354 | (71.5%) | 1.99 |
| Educational Attainment | | | | | $6x10^{-8}$*** |
| Less than 6 years | 208 | (41.7%) | 126 | (25.5%) | |
| 7 – 9 years | 175 | (35.1%) | 192 | (38.8%) | |
| 10 – 12 years | 94 | (18.8%) | 119 | (24.0%) | |
| More than 12 years | 18 | ( 3.6%) | 47 | ( 9.5%) | |
| Body Mass Index (10 years ago) | | | | | 0.18 |
| Underweight | 51 | (10.2%) | 54 | (10.9%) | |
| Normal | 370 | (74.1%) | 348 | (70.3%) | |
| Overweight | 72 | (14.4%) | 78 | (15.8%) | |
| Obese | 6 | ( 1.2%) | 15 | ( 3.0%) | |
| Home Community | | | | | 0.0009*** |
| Cangwu | 125 | (25.1%) | 89 | (18.0%) | |
| Cenxi | 168 | (33.7%) | 160 | (32.3%) | |
| Tengxian | 97 | (19.4%) | 146 | (29.5%) | |
| Wuzhou | 109 | (21.8%) | 100 | (20.2%) | |
| Tobacco Use | | | | | 0.02* |
| Never | 238 | (47.7%) | 273 | (55.2%) | |
| Current | 261 | (52.3%) | 222 | (44.8%) | |
| History of Alcohol Use | | | | | 0.34 |
| Never | 339 | (67.9%) | 357 | (72.1%) | |
| Former | 20 | ( 4.0%) | 15 | ( 3.0%) | |
| Current | 139 | (27.9%) | 123 | (24.8%) | |
| Tea Consumption | 145 | (29.1%) | 155 | (31.3%) | 0.48 |
| Frequency of Salted Fish Consumption in adulthood | | | | | 0.77 |
| Never | 193 | (38.7%) | 197 | (39.8%) | |
| Yearly | 207 | (41.5%) | 202 | (40.8%) | |
| Monthly | 95 | (19.0%) | 85 | (17.2%) | |
| Frequency of Salted Fish Consumption in childhood | | | | | 0.60 |
| Never | 143 | (28.7%) | 146 | (29.5%) | |
| Yearly | 210 | (42.1%) | 213 | (43.0%) | |
| Monthly | 142 | (28.5%) | 125 | (25.3%) | |
| Serum EBV Status | | | | | $1x10^{-15}$*** |
| Negative | 16 | ( 3.2%) | 346 | (69.9%) | |
| Positive | 416 | (83.4%) | 110 | (22.2%) | |
| Unknown | 67 | (13.4%) | 39 | ( 7.9%) | |
| Family History of NPC | | | | | $6x10^{-5}$*** |
| No | 432 | (86.6%) | 473 | (95.6%) | |
| Yes | 54 | (10.8%) | 20 | ( 4.0%) | |
| Missing or Unknown | 13 | ( 2.6%) | 2 | ( 0.4%) | |
| History of Rhinitis | 18 | ( 3.6%) | 22 | ( 4.4%) | 0.61 |
| History of Pharyngitis | 20 | ( 4.0%) | 19 | ( 3.8%) | 0.98 |
| Number of Missing or Repaired Teeth | | | | | 0.30 |
| 0 | 217 | (43.5%) | 202 | (40.8%) | |
| 1 | 63 | (12.6%) | 57 | (11.5%) | |
| 2 | 53 | (10.6%) | 66 | (13.3%) | |
| 3-5 | 83 | (16.6%) | 100 | (20.2%) | |
| 6+ | 83 | (16.6%) | 70 | (14.1%) | |
| Tooth Brushing Frequency | | | | | $6x10^{-11}$*** |
| <=1/day | 325 | (65.1%) | 219 | (44.2%) | |
| >=2/day | 174 | (34.9%) | 276 | (55.8%) | |

†p-value Welch's t-test
§p-value Chi-square test
* $p < 0.05$; ** $p < 0.01$; *** $p < 0.001$

**Table S2. Association between clinical covariates and alpha diversity metrics**

| | **Phylogenetic Diversity** | | **Observed ASVs** | | **Shannon Diversity** | |
|---|---|---|---|---|---|---|
| | statistic | p-value | statistic | p-value | statistic | p-value |
| NPC status | 118.7 | $< 1x10^{-12}$* | 172.6 | $< 1x10^{-12}$* | 46.3 | $1x10^{-11}$* |
| Residential Community | 11.1 | 0.01* | 13.3 | 0.004* | 16.5 | 0.001* |
| Tooth Brushing Frequency | 5.4 | 0.02* | 10.9 | 0.001* | 3.1 | 0.08 |
| History of Pharyngitis | 7.3 | 0.007* | 7.2 | 0.007* | 0.3 | 0.60 |
| Tea Consumption | 1.5 | 0.22 | 0.7 | 0.40 | 5.7 | 0.02* |
| Smoking Status | 0.7 | 0.42 | 0.3 | 0.58 | 4.5 | 0.04* |
| History of Rhinitis | 1.6 | 0.20 | 1.5 | 0.22 | 4.3 | 0.04* |
| Educational Attainment | 4.9 | 0.30 | 5.1 | 0.28 | 7.4 | 0.12 |
| BMI 10 years ago | 3.0 | 0.39 | 4.5 | 0.21 | 1.9 | 0.59 |
| Childhood Salted Fish Consumption | 1.9 | 0.38 | 1.2 | 0.54 | 0.2 | 0.90 |
| Family History | 0.2 | 0.67 | 0.1 | 0.78 | 0.6 | 0.45 |
| Missing or Repaired Teeth | 2.5 | 0.64 | 3.7 | 0.45 | 3.7 | 0.45 |
| Adult Salted Fish Consumption | 1.5 | 0.47 | 1.3 | 0.58 | 0.3 | 0.84 |
| History of Alcohol Use | 0.2 | 0.66 | 0.3 | 0.60 | 0.0 | 0.96 |

*Significant at $\alpha < 0.05$

**Table S3. Model selection for regression against Faith's Phylogenetic Diversity**

| | Model | Adjusted R2 | AIC | ΔAIC | Reference Model |
|---|---|---|---|---|---|
| 1 | host_age + sex + run_num | 0.041 | 4005 | | -- |
| 2 | host_age + sex + run_num + case_control | 0.155 | 3881 | -124 | 1 |
| 3 | host_age + sex + run_num + teeth_brushing_freq | 0.047 | 4000 | -5 | 1 |
| 4 | host_age + sex + run_num + pharyngitis | 0.046 | 4002 | -4 | 1 |
| 5 | host_age + sex + run_num + region_home | 0.046 | 4004 | -2 | 1 |
| 6 | host_age + sex + run_num + case_control + region_home | 0.162 | 3876 | -6 | 2 |
| 7 | host_age + sex + run_num + case_control + pharyngitis | 0.159 | 3877 | -6 | 2 |
| 8 | host_age + sex + run_num + case_control + teeth_brushing_freq | 0.154 | 3883 | 2 | 2 |
| 9 | host_age + sex + run_num + case_control + region_home + pharyngitis | 0.167 | 3871 | -4 | 6 |

**Table S4. Model selection for OObserved ASVs**

| | Model | Adjusted $R^2$ | AIC | ΔAIC | Reference Model |
|---|---|---|---|---|---|
| 1 | host_age + sex + run_num | 0.047 | 10161 | -- | -- |
| 2 | host_age + sex + run_num + case_control | 0.190 | 10000 | -160 | 1 |
| 3 | host_age + sex + run_num + teeth_brushing_freq | 0.057 | 10151 | -10 | 1 |
| 4 | host_age + sex + run_num + region_home | 0.054 | 10157 | -4 | 1 |
| 5 | host_age + sex + run_num + pharyngitis | 0.051 | 10158 | -3 | 1 |
| 6 | host_age + sex + run_num + case_control + region_home | 0.201 | 9989 | -11 | 2 |
| 7 | host_age + sex + run_num + case_control + pharyngitis | 0.194 | 9997 | -4 | 2 |
| 8 | host_age + sex + run_num + case_control + teeth_brushing_freq | 0.189 | 10002 | 1 | 2 |
| 9 | host_age + sex + run_num + case_control + region_home + pharyngitis | 0.205 | 9986 | -3 | 6 |

**Table S5. Model selection for regression against Shannon diversity**

| | Model | Adjusted $R^2$ | AIC | ΔAIC | Reference Model |
|---|---|---|---|---|---|
| 1 | host_age + sex + run_num | 0.036 | 1880 | -- | -- |
| 2 | host_age + sex + run_num + case_control | 0.079 | 1836 | -44 | 1 |
| 3 | host_age + sex + run_num + region_home | 0.048 | 1870 | -10 | 1 |
| 4 | host_age + sex + run_num + tea_consumption | 0.042 | 1875 | -5 | 1 |
| 5 | host_age + sex + run_num + rhinitis | 0.040 | 1877 | -3 | 1 |
| 6 | host_age + sex + run_num + smoker | 0.037 | 1880 | -0 | 1 |
| 7 | host_age + sex + run_num + case_control + region_home | 0.095 | 1821 | -15 | 2 |
| 8 | host_age + sex + run_num + case_control + tea_consumption | 0.084 | 1831 | -5 | 2 |
| 9 | host_age + sex + run_num + case_control + rhinitis | 0.084 | 1832 | -4 | 2 |
| 10 | host_age + sex + run_num + case_control + smoker | 0.082 | 1833 | -3 | 2 |
| 11 | host_age + sex + run_num + case_control + region_home + rhinitis | 0.100 | 1816 | -5 | 7 |
| 12 | host_age + sex + run_num + case_control + region_home + tea_consumption | 0.100 | 1817 | -4 | 7 |
| 13 | host_age + sex + run_num + case_control + region_home + smoker | 0.098 | 1818 | -3 | 7 |
| 14 | host_age + sex + run_num + case_control + region_home + rhinitis + tea_consumption | 0.104 | 1812 | -4 | 11 |
| 15 | host_age + sex + run_num + case_control + region_home + rhinitis + smoker | 0.103 | 1815 | -2 | 11 |
| 16 | host_age + sex + run_num + case_control + region_home + rhinitis + tea_consumption + smoker | 0.106 | 1812 | -6 | 14 |

**Table S8. Species-level identification of co-occurring organism**

| Cluster | Node | Clone Name | HOMD-ID | Status | Identity (%) |
|---|---|---|---|---|---|
| 1 | cPre-1412 | Alloprevotella sp. HMT 308 | HMT-308 | Phylotype | 99.8 |
| | Gran-7770 | Granulicatella adiacens | HMT-534 | Named | 100 |
| | Prev-4b21 | Prevotella histicola | HMT-298 | Named | 99.8 |
| | Prev-b7f2 | Prevotella melaninogenica | HMT-469 | Named | 100 |
| | Prev-cefa | Prevotella salivae | HMT-307 | Named | 100 |
| | Bull-4893 | Solobacterium moorei | HMT-678 | Named | 99.5 |
| | Stre-900d | Streptococcus parasanguinis clade 411 | HMT-411 | Named | 100 |
| | Stre-b566 | Streptococcus salivarius | HMT-755 | Named | 100 |
| | Veil-8f88 | Veillonella atypica | HMT-524 | Named | 100 |
| | Veil-98e7 | Veillonella atypica | HMT-524 | Named | 99.8 |
| | Veil-675d | Veillonella dispar | HMT-160 | Named | 100 |
| 2 | cWee-6b1b | Bergeyella sp. HMT 322 | HMT-322 | Phylotype | 99.8 |
| | Gran-5a37 | Granulicatella adiacens | HMT-534 | Named | 99.8 |
| | Haem-7aaf | Haemophilus parainfluenzae | HMT-718 | Named | 100 |
| | Pept-facb | Peptococcus sp. HMT 168 | HMT-168 | Phylotype | 99.5 |
| | Porp-fd4f | Porphyromonas pasteri | HMT-279 | Named | 100 |
| | Prev-fb41 | Prevotella aurantiaca | HMT-943 | Named | 99.8 |
| | Prev-71e7 | Prevotella melaninogenica | HMT-469 | Named | 99.8 |
| | Prev-04b7 | Prevotella nanceiensis | HMT-299 | Named | 99.8 |
| | Prev-603e | Prevotella shahii | HMT-795 | Named | 100 |
| | Stre-0531 | Streptococcus parasanguinis clade 411 | HMT-411 | Named | 99.5 |
| | Veil-6e24 | Veillonella parvula | HMT-161 | Named | 99.5 |

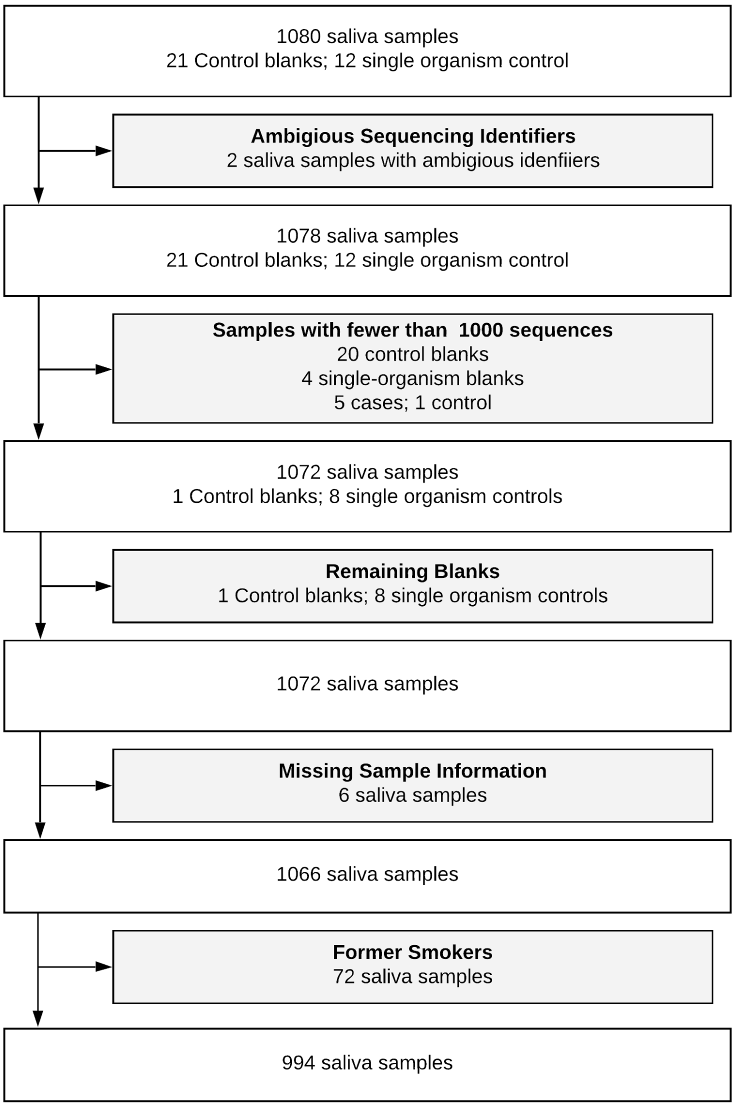


**Figure S1. Flowchart for sample selection**

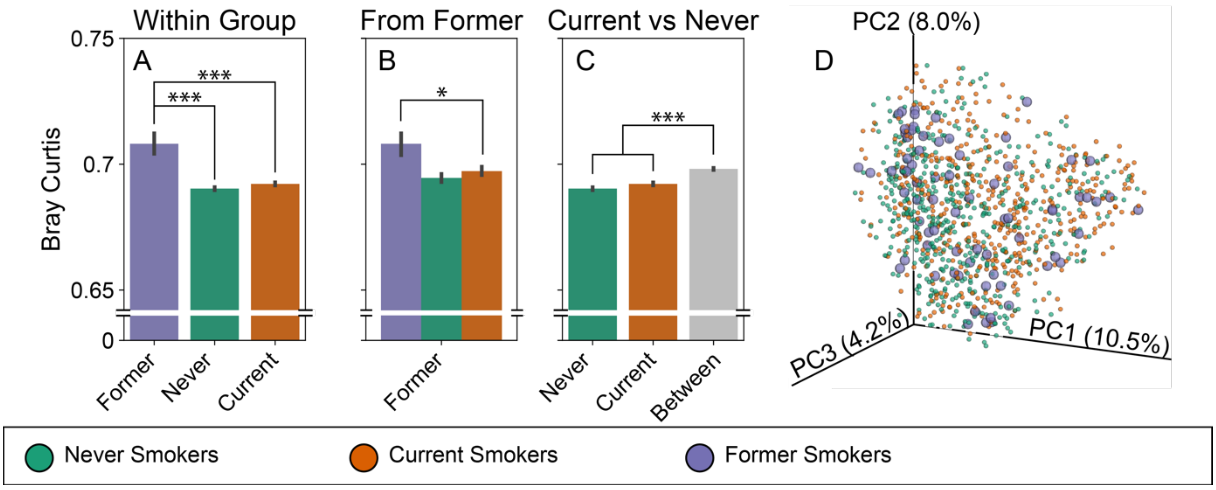


**Figure S2. Former Smokers have highly heterogenous microbiomes.** Bray Curtis distance highlights the high degree of heterogeneity among former smokers. The distribution of distances (A-C) shows the mean distance (± bootstrapped 95% CI) between the group labeled on the bottom and the group described by the color. (A) The within-group distance for former smokers (purple) is greater than the within-group distance for either never (teal) or current (orange) smokers, despite the larger group sizes. Significance is calculated using a permutive pairwise t-test. (B) The within-group distance for former smokers (purple) is greater than the distance from former smokers to never smokers (teal) or to current smokers (orange); significance from pairwise-permanova. (C) The distance between never and current smokers (gray) is greater than the within group distance for either group; this difference is significantly different in a pairwise Permanova. (D) PCoA projection of Bray-Curtis highlighting the Former smokers (large spheres). *** Permutative $p = 0.001$; ** $p < 0.01$; * $p < 0.05$; n=999 permutations for all tests.

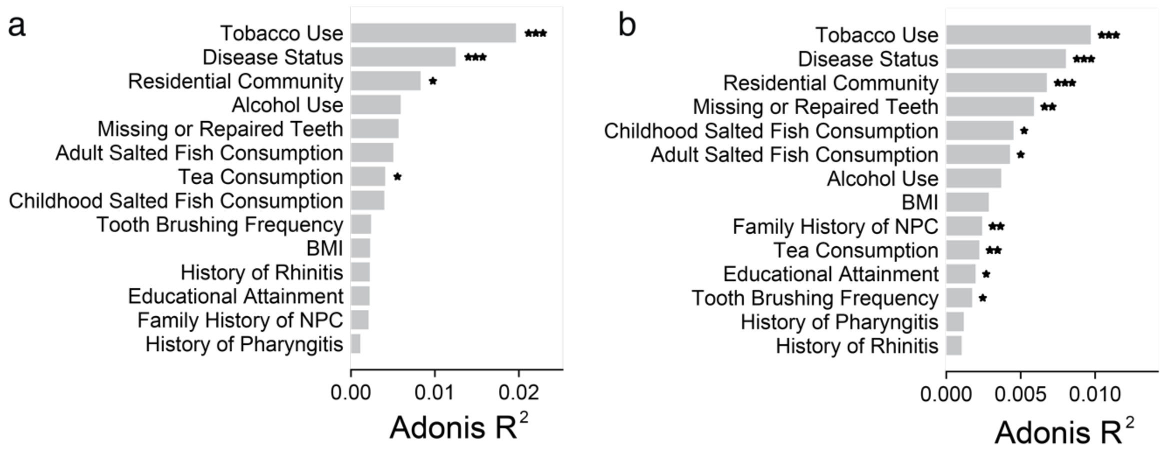


**Figure S3 A community of abundant ASVs retains the difference in the microbiome between NPC cases and controls.** Univariate Adonis testing with a model adjusted for age, sex, and sequencing run in (**a**) weighted UniFrac distance and (**b**) Bray Curtis; tobacco use has more explanatory power than disease status. Significance is indicated as *** FDR adjusted p < 0.001; ** p < 0.01; * p < 0.05.

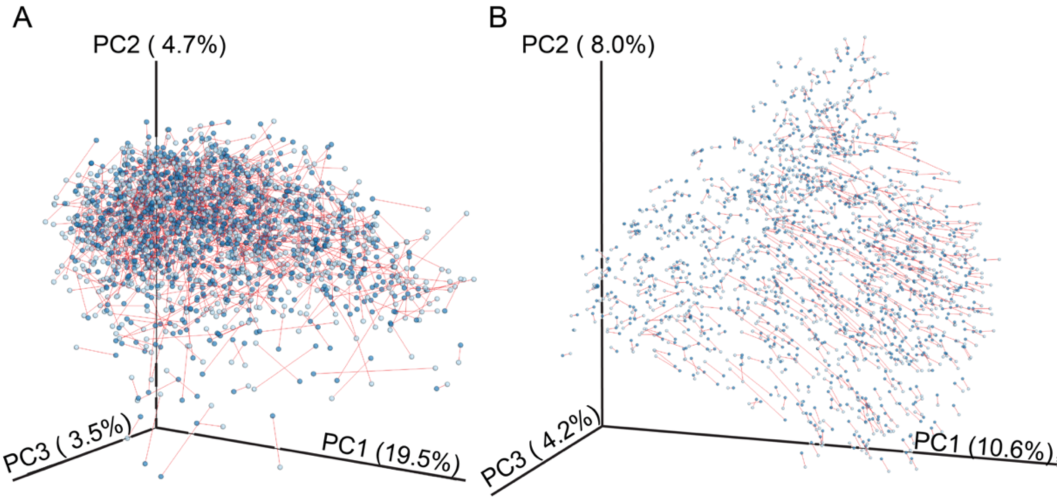


**Figure S4. The abundant ASVs recapitulate community structure well.** Procrustes analysis comparing filtered taxa (light blue) to the full set (dark blue) using (A) unweighted UniFrac and (B) Bray Curtis distance. Paired samples are linked by red lines. Both sets of data are significantly correlated (mantel p = 0.001**,** 999 permutations). The correlation coefficient for Bray-Curtis distance is 0.96 and the correlation for unweighted UniFrac distance is 0.75.

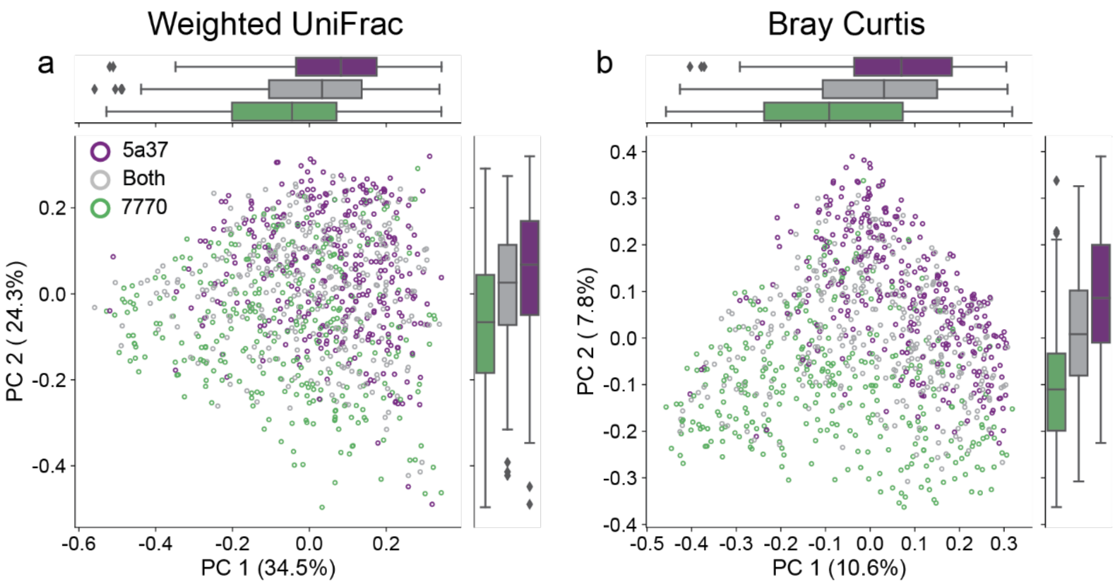


Figure S5. ***Granulicatella adiacens* variant predicts community structure**. PCoA for (a) weighted UniFrac distance and (b) Bray-Curtis distance shows the G. *adiacens* variant (Gran-5a37 alone, purple; Gran-7770 alone, green; or both, gray), shows separation about PC1 and PC2. The *G. adiacens* variant is significantly associated with differences in beta diversity in both metrics (p = 0.001, 999 permutations)
